# Visualization and analysis of single-cell RNA-seq data by kernel-based similarity learning

**DOI:** 10.1101/052225

**Authors:** Bo Wang, Junjie Zhu, Emma Pierson, Daniele Ramazzotti, Serafim Batzoglou

**Affiliations:** Department of Computer Science, Stanford University; Department of Electrical Engineering, Stanford University; Department of Pathology, Stanford University

**Keywords:** single-cell analysis, similarity learning, dimension reduction, data visualization

## Abstract

Single-cell RNA-seq technologies enable high throughput gene expression measurement of individual cells, and allow the discovery of heterogeneity within cell populations. Measurement of cell-to-cell gene expression similarity is critical to identification, visualization and analysis of cell populations. However, single-cell data introduce challenges to conventional measures of gene expression similarity because of the high level of noise, outliers and dropouts. Here, we propose a novel similarity-learning framework, SIMLR (single-cell interpretation via multi-kernel learning), which learns an appropriate distance metric from the data for dimension reduction, clustering and visualization applications. Benchmarking against state-of-the-art methods for these applications, we used SIMLR to re-analyse seven representative single-cell data sets, including high-throughput droplet-based data sets with tens of thousands of cells. We show that SIMLR greatly improves clustering sensitivity and accuracy, as well as the visualization and interpretability of the data.

## Background

Single-cell RNA sequencing (scRNA-seq) technologies have recently emerged as a powerful means to measure gene expression levels of individual cells and to reveal previously unknown heterogeneity and functional diversity among cell populations [1]. Quantifying the variation across gene expression profiles of individual cells is key to the identification and analysis of complex cell populations that arise in neurology [2], immunology [3], oncology [4] and developmental biology [5]. The heterogeneity identified across individual cells can answer questions irresolvable by traditional ensemble-based methods, where gene expression measurements are averaged over a population of cells pooled together [6], [7]. Recent studies have demonstrated that *de novo* cell type discovery and identification of functionally distinct cell subpopulations are possible via unbiased analysis of all transcriptomic information provided by scRNA-seq data [8]. Therefore, unsupervised clustering of individual cells using scRNA-seq data is critical to developing new biological insights and validating prior knowledge.

Many existing single-cell studies employ computational and statistical methods that have been developed primarily for analysis of data from traditional bulk RNA-seq methods [8]–[10]. These methods do not address the unique characteristics that make single-cell expression data especially challenging to analyze: outlier cell populations, transcript amplification noise, and biological effects such as the cell cycle [11]. In addition, it has been shown that many statistical methods fail to alleviate other underlying challenges, such as dropout events, where zero expression measurements occur due to sampling or stochastic transcriptional activities [12]. Recently, new single-cell platforms such as DropSeq [13], InDrop [14] and GemCode single-cell technology [15] have enabled a dramatic increase in throughput to thousands of cells. These platforms have adapted recent sequencing protocols such as unique molecular identifiers (UMIs) to create digital counts of transcripts in a cell. However, with 3'-end sequencing instead of full transcript sequencing, low coverage per cell and varying capture efficiency, these high-throughput platforms produce data sets where 95% of measurements are zeros.

The technological differences across single-cell platforms as well as the biological differences across studies can reduce the usability of unsupervised clustering methods. Core to the problem is that unsupervised clustering methods usually rely on specific similarity metrics across the objects to be clustered; standard similarity metrics may not generalize well across platforms and biological experiments, and thus be unsuitable for scRNA-seq studies. To address this problem and answer the key question of “which cells are similar or different” in a way that generalizes across different single-cell data sets, we introduce SIMLR (single-cell interpretation via multi-kernel learning), a novel framework that learns an appropriate cell-to-cell similarity function from the input single-cell data. SIMLR simultaneously clusters cells into groups for subpopulation identification and produces a 2-D or 3-D visualization of the expression data, with the same similarity function applied for clustering and low-dimensional projection. As a result, the identified cell subpopulations and their separation are intuitively visualized.

SIMLR offers three main unique advantages over previous methods. First, it *learns* a distance metric that best fits the structure of the data via combining multiple kernels. This is important because the diverse statistical characteristics due to varying noise and dropout patterns in single-cell data produced today do not easily fit specific statistical assumptions made by standard dimension reduction algorithms. The adoption of multiple kernel representations allows greater flexibility in defining cell-to-cell similarities than a single kernel or similarity measures such as correlation and Euclidian distance. Second, SIMLR addresses the challenge of high levels of dropout events that can significantly weaken cell-to-cell similarities even under an appropriate distance metric, by employing graph diffusion [16], which improves weak similarity measures that are likely to result from noise or dropout events. Third, in contrast to some previous analyses that pre-select gene subsets of known function [10], [17], SIMLR is unsupervised, thus allowing *de novo* discovery from the data. We empirically demonstrate that SIMLR produces more reliable clusters than commonly used linear methods, such as principal component analysis (PCA) [18], and nonlinear methods, such as t-distributed stochastic neighbor embedding (t-SNE) [19], and we use SIMLR to provide 2-D and 3-D visualizations that assist with the interpretation of single-cell data derived from several diverse technologies and biological samples.

## Results

### Overview of Algorithm

Here we highlight the main ideas in our methodology underlying SIMLR, and we provide full details in **Materials and Methods**.

Given a *N*×*M* gene expression matrix *X* with *N* cells and *M* genes (*N* < *M*) as an input, SIMLR solves for *S*, a *N*×*N* symmetric matrix that captures pairwise similarities of cells. In particular, *S_ij_*, the (*i, j*)-th entry of *S*, represents the similarity between cell *i* and cell *j*. SIMLR assumes that if *C* separable populations exist among the *N* cells, then *S* should have an approximate block-diagonal structure with *C* blocks whereby cells have larger similarities to other cells within the same subpopulations.

We introduce an optimization framework that learns *S* by incorporating multiple kernels to learn appropriate cell-to-cell distances from the data (Figure 1a) where each kernel corresponds to a simple distance measure. We provide an efficient algorithm to optimize for *S*, while simultaneously learning the block-diagonal structure within !. The cell-to-cell similarity values in *S* can be used to create an embedding of the data in 2-D or 3-D for visualization, as well as a projection of the data into a latent space of arbitrary dimension to further identify groups of cells that are similar (Figure 1b).

**Figure 1.**
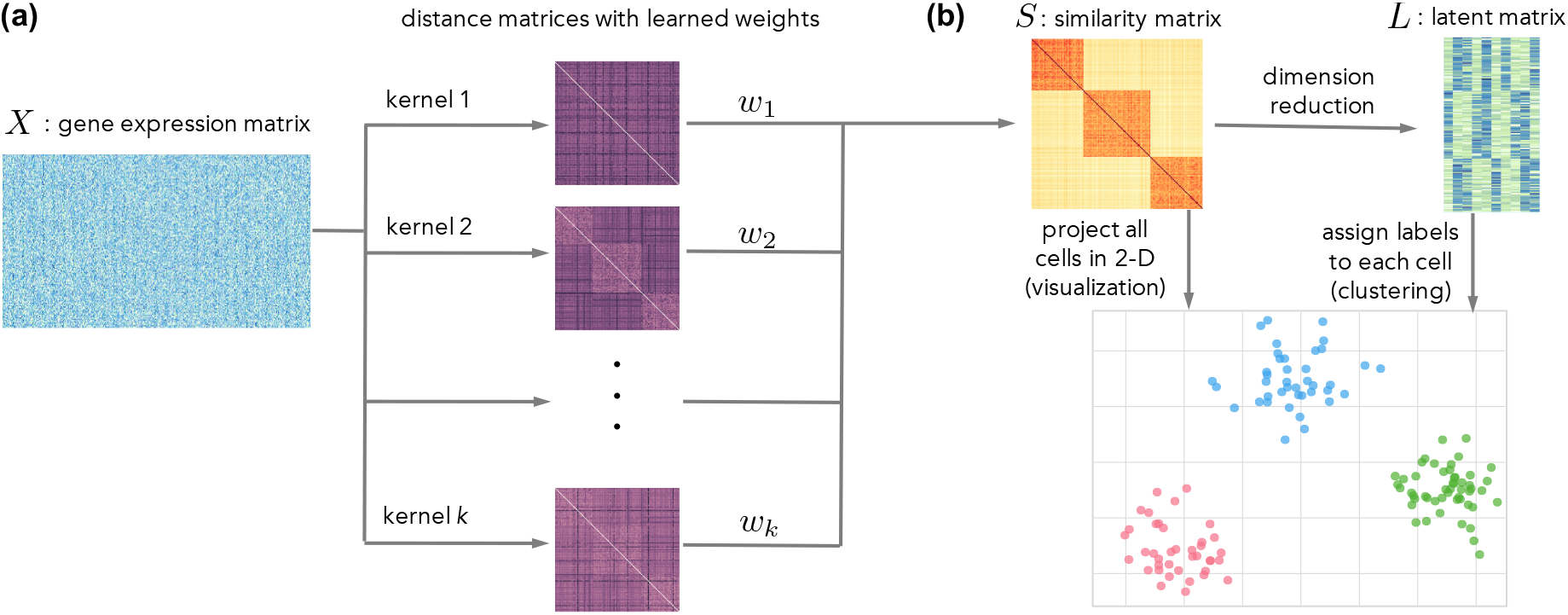
Outline of S1MLR. **(a)** SIMLR learns a proper metric for the cell-to-cell distances using the gene expression and constructs a similarity matrix. **(b)** The similarity matrix is used for visualization of cells in 2-D and for dimension reduction for clustering.

#### General optimization framework

SIMLR computes cell-to-cell similarities through the following optimization framework:

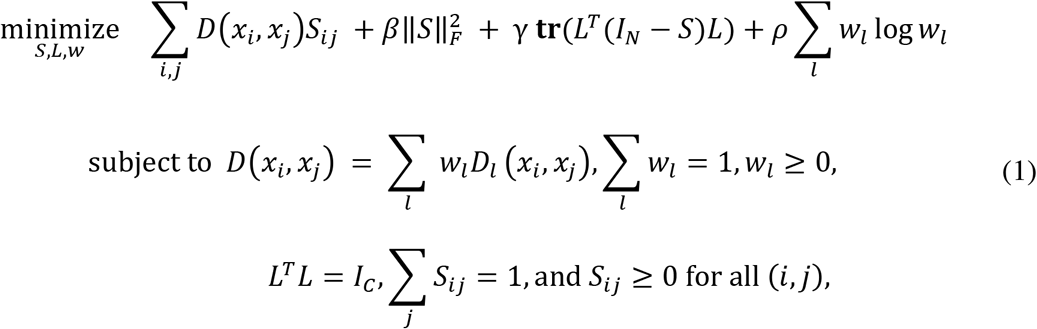

where *x_i_* is the length-*M* gene expression vector of cell *i*, i.e., the *i*th row of *X*; *D*(*x_i_*, *x_j_*) is the distance between cell *i* and cell *j*, expressed as a linear combination of distance metrics *D_l_* with weights *w_l_*; *I_N_* and *I_C_* are *N*×*N* and *C*×*C* identity matrices respectively, and *β* and *γ* are non-negative tuning parameters. ‖*S*‖*_F_* denotes the Frobenius norm of *S*, and *L* denotes an auxiliary low-dimensional matrix enforcing the low rank constraint on *S*. The optimization problem involves solving for three variables: the similarity matrix *S*, the weight vector *w*, and a *N*×*C* rank-enforcing matrix *L*.

The intuition behind the first term in the formula is that the learned similarity *S* between two cells should be small if the distance between them is large. The second term is a regularization term that prevents the learned similarities from becoming too close to an identity matrix (**Supplementary Figure 27**). If there are *C* subpopulations, the gene expressions of cells of the same sub-type should have high similarity, and ideally the effective rank of *S* should be *C*. Thus, the third term along with the constraint on *L* enforces the low-rank structure of *S*: the matrix (*I_N_* − *S*) is essentially the graph Laplacian [20], and the trace-minimization problem enforces approximately *C* connected components in a similarity graph that consists of nodes representing the cells, and edge weights corresponding to pairwise similarity values in *S* [20]. The fourth term imposes constraints on the kernel weights to avoid selection of a single kernel; we empirically found that this regularization improves the quality of learned similarity (**Supplementary Table 1**).

One critical component of this optimization problem is the choice of the distance measure *D*(*x_i_*, *x_j_*) between pairs of cells. It is well-known that the distance metric defined for the input space is critical to the performance of clustering and visualization algorithms designed for high-dimensional data [21]. Due to the presence of outliers and unusual zero-inflated distributions in single-cell data, standard metrics like the Euclidian distance may fail to perform well. Thus, instead of using a pre-defined distance metric, we incorporate multiple kernel learning [22] that flexibly combines multiple distance metrics. Multiple kernels have been shown to correspond to different informative representations of the data and often are superior to single kernels in various machine learning contexts [23], [24].

Even though this general optimization problem is non-convex, we are able to employ a computationally efficient algorithm for optimizing *S*, *L* and *w* in **Algorithm 1** (with full details for each step in **Materials and Methods)**. The intuition behind our procedure is simple: holding two of these three variables fixed, the optimization problem over the third variable is convex. Hence, we alternate between optimizing each variable while holding the other two fixed until convergence. Step 4 in **Algorithm 1** is an auxiliary step where we incorporate a similarity enhancement heuristic based on the graph diffusion process (with full details in **Materials and Methods**). The intuition behind diffusion-based similarity enhancement is two-fold: (1) diffusion processes adds “transitive” similarities between two seemingly dissimilar cells that also have many common neighboring cells with high similarity. This enables SIMLR to alleviate the impact of dropout events and noise typical in scRNA-seq data. (2) The similarity matrix *S* obtained from the optimization framework may contain arbitrarily weakened entries due to the constraints on *S*, so higher order structures such as local connectivity can be exploited via the diffusion process to improve the numerical stability of the solution.

#### Dimension reduction for clustering and visualization

SIMLR relies on the stochastic neighbor embedding (SNE) [25] methodology for dimension reduction, with an important modification: t-SNE computes the similarity of the high-dimensional data points using a Gaussian kernel as a distance measure and projects the data onto a lower dimension that preserves this similarity. Instead of using the gene expression matrix as an input to t-SNE, we use the learned cell-to-cell similarity *S* (detailed in **Supplementary Note 3.9**).

For visualization, we use our modified t-SNE algorithm to project the data into two or three dimensions so that the hidden structures in the data can be depicted intuitively. For clustering, we use the same approach to reduce the dimensions to B, resulting in an *N*×*B* latent matrix *Z*, to which we can apply any existing clustering algorithms, such as K-means [26] to assign labels to each cell. The number of reduced dimensions *B* is by default equal to the number of desired clusters *C*.

#### Gene prioritization using the learned similarity

The learned cell-to-cell similarity matrix *S* can be leveraged to perform gene prioritization: we rank each gene by measuring how its expression values across the cells correlate with the learned cell-to-cell similarity.

Given the similarity *S* and the expression of a gene across all cells, *f* we adopt the Laplacian score,

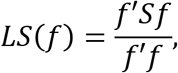

a well-known unsupervised feature ranking method [27] to measure the concordance between genes and similarity. A higher the Laplacian score, the more important the gene is to globally differentiate the subpopulations of cells. However, the Laplacian score is typically very sensitive to noise in the measured similarity. To overcome this issue, we propose a bootstrap approach for which we randomly subsample a proportion of cells (e.g., 80% of the total number of cells), and rank genes based on the Laplacian score on the subset of learned similarity. We aggregate rankings from independent trials using Robust Ranking Aggregation (RRA) [28] to finalize the prioritized genes (**Supplementary Note 3.13**). This final ranking reflects the importance of each gene to the similarity learned by SIMLR. We also empirically show that our gene prioritization based on SIMLR can reveal biologically meaningful genes that define cell subpopulations without any prior information (**Supplementary Figures 14-15, Supplementary Note 3.2**).

#### Extension of SIMLR's framework for very large-scale datasets

Recent developments in scRNA-seq technologies have led to massively parallel profiling of tens of thousands of single cells in one experiment. To address this computational challenge, we provide an additional fast approximate implementation as an extension of SIMLR's framework. Instead of using a dense similarity matrix, we employ a sparse similarity setting where only the top nearest neighbors of each cell are allowed to have nonzero similarities. The sparse similarity framework not only facilitates the calculation of the top eigenvectors of the Laplacian matrix when solving for *L* in Eqn. (1), but also reduces the memory complexity of SIMLR. A significant speed-up results from limiting each similarity update to involve only the top k nearest neighbors of each cell (**Supplementary Note 3.12.1**). Moreover, the scalability of downstreaming applications of SIMLR such as clustering (**Supplementary Note 3.12.2**) and visualization (**Supplementary Note 3.12.3**) are also improved in this framework. Results on specific large-scale scRNA-seq datasets are described in the following sections.

## Applications

### Learning cell-to-cell similarities on data sets with ground-truth

We start by benchmarking SIMLR against conventional predefined measures in capturing true cell-to-cell similarities on four published single-cell data sets (for the full details of each data set, see **Materials and Methods**):

1. Eleven cell populations including neural cells and blood cells (Pollen data set [9]).
2. Neuronal cells with sensory subtypes (Usoskin data set [8]).
3. Embryonic stem cells under different cell cycle stages (Buettner data set [17]).
4. Pluripotent cells under different environment conditions (Kolodziejczk data set [10]).

We selected these data sets because they span a variety of cell types and have different numbers of subpopulations, representing a wide range of single-cell data; cell types in each data set were known *a priori* and were further validated in the respective studies, providing a reliable gold standard with which to assess clustering performance (Table 1). To evaluate SIMLR's performance on these data sets, we compared subpopulation labels assigned after dimension reduction with SIMLR to the true subpopulation labels from the respective studies.

**Table 1.** Summary of the characteristics of the four real single-cell data sets, with SIMLR's computational time.

| Data set | # cells | # genes | # populations | run time (seconds) |
| --- | --- | --- | --- | --- |
| Pollen [9] | 249 | 14805 | 11 | 3.40 |
| Usoskin [8] | 622 | 17772 | 4 | 18.54 |
| Buettner [17] | 182 | 8989 | 3 | 2.369 |
| Kolodziejczyk [10] | 704 | 10685 | 3 | 23.67 |

SIMLR was given the raw gene expressions and the true number of clusters, but no information about the true labels. To assess how well the learned similarities agree with the validated cell identities (i.e., true labels), we visualize the cell-to-cell similarities by ordering the cells by their true labels so that cells with the same label are grouped together. We compared the cell-to-cell similarities learned by SIMLR with a similarity matrix computed from Gaussian kernels applied to Euclidean distances (Euclidean Similarity), and a pairwise correlation matrix (Pearson Correlation) (Figure 2). We plotted the resulting symmetric similarity matrices, with colors indicating the known labels, and observed that SIMLR learned a similarity matrix with block structures in remarkable agreement with the previously validated labels, while the other similarity matrices agree less obviously (Figure 2). Both Euclidean distances and Pearson correlations are sensitive to outliers, and Pearson correlations do not capture nonlinear relationships, so spurious similarities between cells from different groups surface with these predefined measures. On the other hand, SIMLR handles outliers with its rank constraint and nonlinear similarities using multiple kernels, making its learned similarity more suitable for single-cell data sets.

**Figure 2.**
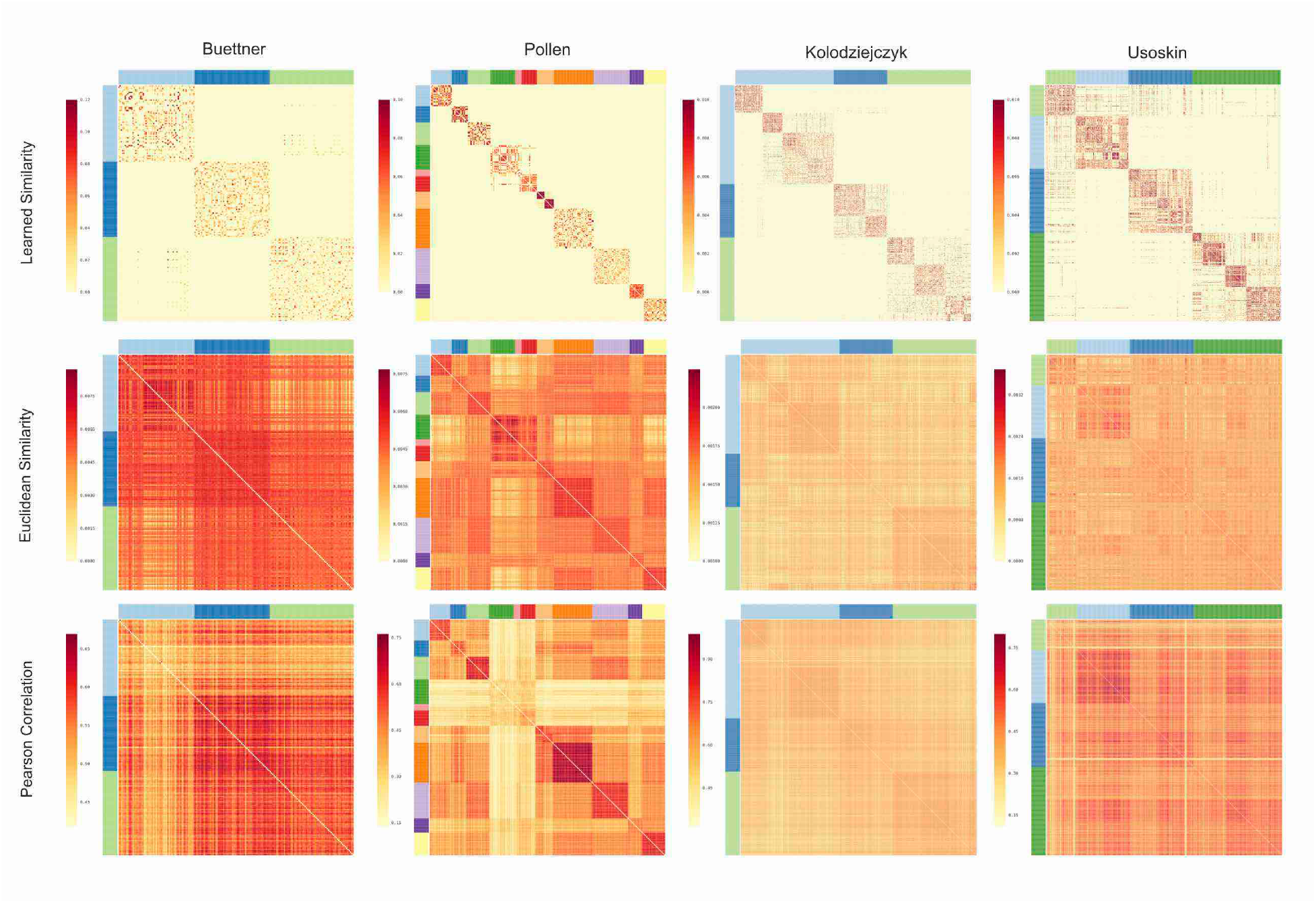
Heatmaps of similarities of cells in four data sets. Three types of similarities are compared: (1) similarities learned from the data by SIMLR, (2) similarities computed from Gaussian kernels applied to Euclidean distances, and (3) similarities computed from pairwise Pearson correlations.

Because true labels of the Buettner data set are cell-cycle states that have been well-studied in the literature, we conducted additional analysis on this data set using SIMLR's gene prioritization. We found that SIMLR is able to reveal strongly connected interaction networks on the prioritized genes by querying the STRING database [29]. When partitioning these genes based on cell-state-specificity, we were also able to identify a large number of highly significant biological processes related to cell-cycle, translation and metabolic processes (**Supplementary Figures 14-15, Supplementary Note 3.7.1**).

Furthermore, SIMLR demonstrates one additional advantage: it can identify additional subpopulation structures even when the number of clusters input into the algorithm is conservatively selected. From the similarity structure learned by SIMLR in the Kolodziejczyk data set we observed that each of the three validated clusters could be further divided into sub-clusters, which is consistent with the unsupervised analysis in the corresponding study [10]. However, these sub-clusters were identified using a small number of pre-selected genes in the original study. SIMLR preserved these substructures in an unbiased fashion while having clearly removed the spurious similarities between cells from different (validated) groups. In the remainder of our analysis, as gold standard we use only the unbiased and validated true labels as opposed to the additional subpopulations identified by unsupervised analysis.

### Reducing the dimension of scRNA-seq data

To analyze well how SIMLR performs dimension reduction, we compare SIMLR's dimension reduction application to other dimension reduction methods by comparing the latent space projection produced by SIMLR to the latent space projections produced by other methods. Following a supervised approach used by [12] to compare different dimension reduction methods, we classify each cell based on the true labels of its nearest neighbors in the dimension-reduced space, and assess the accuracy of this classification using cross validation; we refer to this metric as nearest neighbor error (NNE) (**Supplementary Note 3.1.5**). This test assesses how accurately new cells can be classified using cells whose labels are already known.

In addition to using the NNE, we also compare different dimension reduction results following an unsupervised approach: we use K-means [26] to cluster cells in these different latent spaces and assess which latent-space clustering has the greatest concordance with the true labels. When comparing different dimension reduction methods combined with K-means, we assume the number of clusters *C* is known *a priori.* The performance metric we use to compare the results to the true cell identities is the normalized mutual information (NMI, **Supplementary Note 3.1.1**), a standard measure of clustering concordance [28].

We select these two types of performance metrics for dimension reduction specifically because they measure different aspects of the data in the low dimensional latent space: NNE directly reflects how closely cells from the same population are surrounded by each other, whereas NMI provides a global view of how well cells from different populations are separated. We also tested four other similar performance metrics in **Supplementary Note 3.1**, shown in **Supplementary Figures 2-5**, to ensure that SIMLR performed well under different standards.

We performed extensive comparisons of SIMLR with 8 other dimension reduction methods on the four data sets to test its utility for dimension reduction. The 8 methods included standard linear methods including PCA [18], FA [33], and probabilistic PCA (PPCA) [28]; nonlinear methods including t-SNE [23], Laplacian eigenmaps [25], multidimensional scaling (MDS) [26], and Sammon mapping [27]; and model-based methods specifically designed for single-cell data like zero-inflated factor analysis (ZIFA) [12]. The methods we tested included all methods used in analyses of the original data sets. In addition, the Pollen and the Usoskin data sets were the only ones with validated labels used in [12] to assess ZIFA, which was specifically designed for single-cell data.

The NMI and NNE values for the 9 methods are summarized in Table 2 and Table 3 respectively. SIMLR consistently outperforms the existing alternatives on the four data sets, and most of the differences in NMI and NNE between SIMLR and the second best method are remarkably large.

**Table 2.** NMI values for the four single-cell data sets to compare different dimension methods. Higher values indicate better performance.

| Data set | PCA | Laplacian | MDS | t-SNE | Sammon | PPCA | FA | ZIFA | SIMLR |
| --- | --- | --- | --- | --- | --- | --- | --- | --- | --- |
| **Buettner** | 0.56 | 0.27 | 0.56 | 0.32 | 0.05 | 0.59 | 0.67 | 0.65 | **0.89** |
| **Kolodziejczyk** | 0.77 | 0.62 | 0.77 | 0.77 | 0.40 | 0.76 | 0.72 | 0.72 | **0.99** |
| **Pollen** | 0.83 | 0.80 | 0.83 | 0.93 | 0.75 | 0.83 | 0.82 | 0.79 | **0.95** |
| **Usoskin** | 0.39 | 0.48 | 0.39 | 0.69 | 0.41 | 0.40 | 0.36 | 0.41 | **0.74** |

**Table 3.** NNE values for the four single-cell data sets to compare different dimension methods. Lower values indicate better performance.

| Data set | PCA | Laplacian | MDS | t-SNE | Sammon | PPCA | FA | ZIFA | SIMLR |
| --- | --- | --- | --- | --- | --- | --- | --- | --- | --- |
| Buettner | 0.21 | 0.38 | 0.22 | 0.18 | 0.21 | 0.20 | 0.11 | 0.16 | 0.05 |
| Kolodziejczyk | 0.0016 | 0.0278 | 0.0016 | 0.0014 | 0.018 | 0.0016 | 0.009 | 0.01 | 0.0018 |
| Pollen | 0.052 | 0.156 | 0.055 | 0.023 | 0.11 | 0.056 | 0.075 | 0.075 | 0.020 |
| Usoskin | 0.30 | 0.11 | 0.29 | 0.072 | 0.20 | 0.29 | 0.31 | 0.28 | 0.063 |

To test the robustness of SIMLR, we conducted three additional sensitivity experiments for each data set.

1. We used varying numbers (3 – 20) of latent dimensions *B* to evaluate the performance of SIMLR and other methods. This evaluation is critical because typically the true number of clusters in the data is unknown. (As mentioned before, ideally *B* should be equal to the true number of clusters for SIMLR.)
2. We dropped varying fractions (5% – 70%) of the gene measurements in the input gene expression matrix to analyze how each method performs when random levels of dropout are present across the data, which is relevant to the high dropout rate in single-cell data.
3. We added independent zero-mean Gaussian noise with varying variances *σ*^2^ (0.1 – 1) to the gene expression matrix. The number of dimensions *C* used in this experiment is set to be the true number of clusters. The number of dimensions *C* used in this experiment is set to be the true number of clusters. In order to preserve the dropout characteristics, we ensured that the added noise was set to zero at a frequency equal to the dropout rate in each data set. Formally, we added a random noise vector y_*t*_ to *x_t_* by the following process:

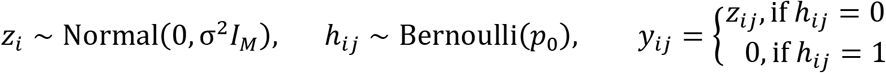

where *p_0_* is the dropout rate (proportion of zeros) in the original expression matrix *X*. We set the values of an entry in the new expression matrix to zero if its value after adding noise dropped to zero or below.

The NMI and NNE values (Figure 3a, b) on the Buettner data set show remarkably better performance by SIMLR as compared to other methods. In addition, we observe that SIMLR is not sensitive to the number of latent dimensions *B* even though the most suitable choice of *B* should ideally be the number of clusters *C* if it is known. ZIFA and FA achieve high NMI values at certain values of *B* but are less stable and can perform much worse than SIMLR otherwise (as is shown in the first column of Figure 3). Moreover, as the fraction of genes increases, the NMI increases and the NNE decreases more noticeably for SIMLR. Finally, while the performance of SIMLR decreases with the noise variance, it still outperforms other methods.

**Figure 3.**
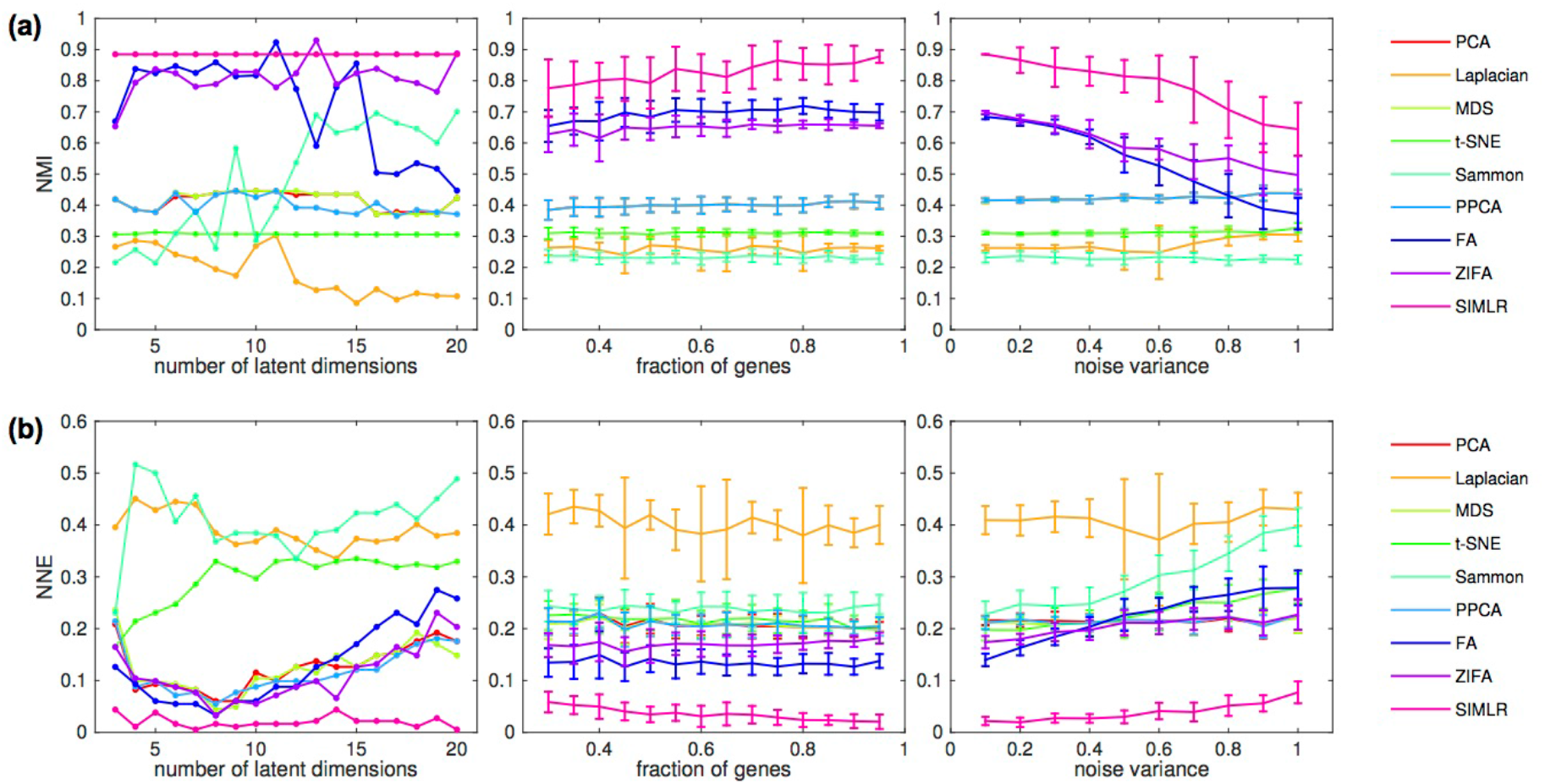
Comparison of 9 different dimension reduction methods under the three robustness experiments on the Buettner data set. **(a)** The clustering accuracy in terms of NMI; higher scores denote better performance. **(b)** The clustering accuracy in terms of NNE; lower scores denote better performance. SIMLR is not sensitive to the number of latent dimensions selected for the algorithm (left column), and still outperforms other methods even when fractions of genes are dropped (middle column) or when additional noise is added to the data (right column).

Based on other metrics, including Clustering Accuracy [30] and Purity [31], on all four data sets (shown in **Supplementary Figures 2-5**), we observe that SIMLR also outperforms other methods on the three sensitivity experiments above and no other method dominates in specific regimes.

### Grouping single cells using clustering algorithms

To assess SIMLR's clustering application in the previous section, we used the following two-stage procedure: (1) reduce the data to a C-dimensional latent space using SIMLR with input parameter *C*, and (2) apply K-means clustering with input number of clusters equal to *C* to compute the cluster assignment of each cell. However, because SIMLR is a similarity framework, it can be flexibly adopted in other clustering frameworks that directly take similarities as inputs. So we also assess SIMLR combined with Affinity Propagation (AP) [32] (rather than k-means).

We found that K-means clustering with SIMLR consistently outperforms the existing alternatives: Non-Negative Matrix Factorization (NMF) [34], Gaussian Mixture Models (GMM) [35], Dirichlet Process Mixture Models (DPMM) [36], on the four data sets (Table 4). These three clustering methods are model-based algorithms that have been used in recent single-cell studies [37], [38]. Further, we conducted extra experiments with AP which takes similarities as inputs to demonstrate that the similarities learned by SIMLR (used for SIMLR+AP) can significantly outperform the other simpler similarities, such as Euclidean similarity (used for Euc.+AP) and Pearson correlation (used for Corr.+AP). This superior clustering performance is expected from plotting of the similarity matrices (Figure 2).

**Table 4.**
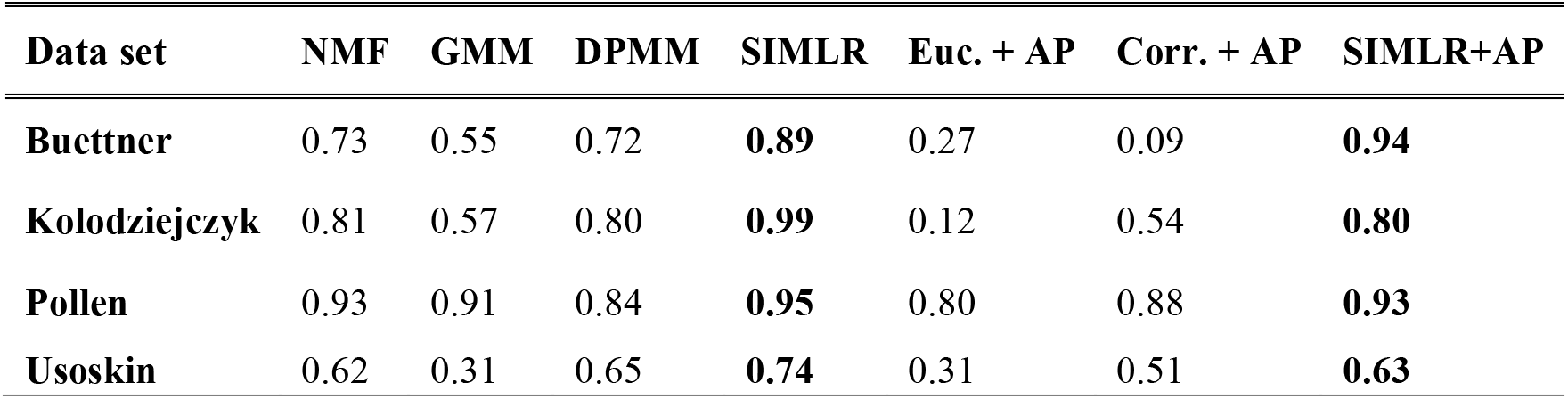
NMI values for the four single-cell data sets to compare different clustering algorithms. Higher values indicate better performance.

| **Data set** | **NMF** | **GMM** | **DPMM** | **SIMLR** | **Euc. + AP** | **Corr. + AP** | **SIMLR+AP** |
| --- | --- | --- | --- | --- | --- | --- | --- |
| **Buettner** | 0.73 | 0.55 | 0.72 | **0.89** | 0.27 | 0.09 | **0.94** |
| **Kolodziejczyk** | 0.81 | 0.57 | 0.80 | **0.99** | 0.12 | 0.54 | **0.80** |
| **Pollen** | 0.93 | 0.91 | 0.84 | **0.95** | 0.80 | 0.88 | **0.93** |
| **Usoskin** | 0.62 | 0.31 | 0.65 | **0.74** | 0.31 | 0.51 | **0.63** |

Clustering methods typically require the number of clusters as an input parameter in order to reveal meaningful grouping in the data. Thus, when applied to clustering, the input parameter *C* to SIMLR can be conveniently assigned to be the number of clusters used as an input for the downstream clustering algorithm. However, if number of clusters is not known a *priori*, SIMLR provides two heuristics based on the structures in the similarities to estimate an optimal choice of *C* (**Supplementary Note 3.10**). Effectiveness of these two heuristics is demonstrated on the four single-cell datasets (**Supplementary Figures 19-20**)

### Visualizing cells in 2-D

After confirming that cell-to-cell similarities learned by SIMLR are meaningful and that the clustering performance of SIMLR is reliable, we applied SIMLR's SNE-based dimension reduction and visualization in 2-D to verify that the structures in the data are visually intuitive. We compared SIMLR with two of the most commonly used visualization methods in single-cell analysis: PCA and t-SNE. We also included ZIFA, which was shown to outperform many other model-based methods [15]. In the resulting visualizations (Figure 4), none of the four methods used the true labels as inputs for dimension reduction, and the true label information was added in the form of distinct colors to validate the results.

**Figure 4.**
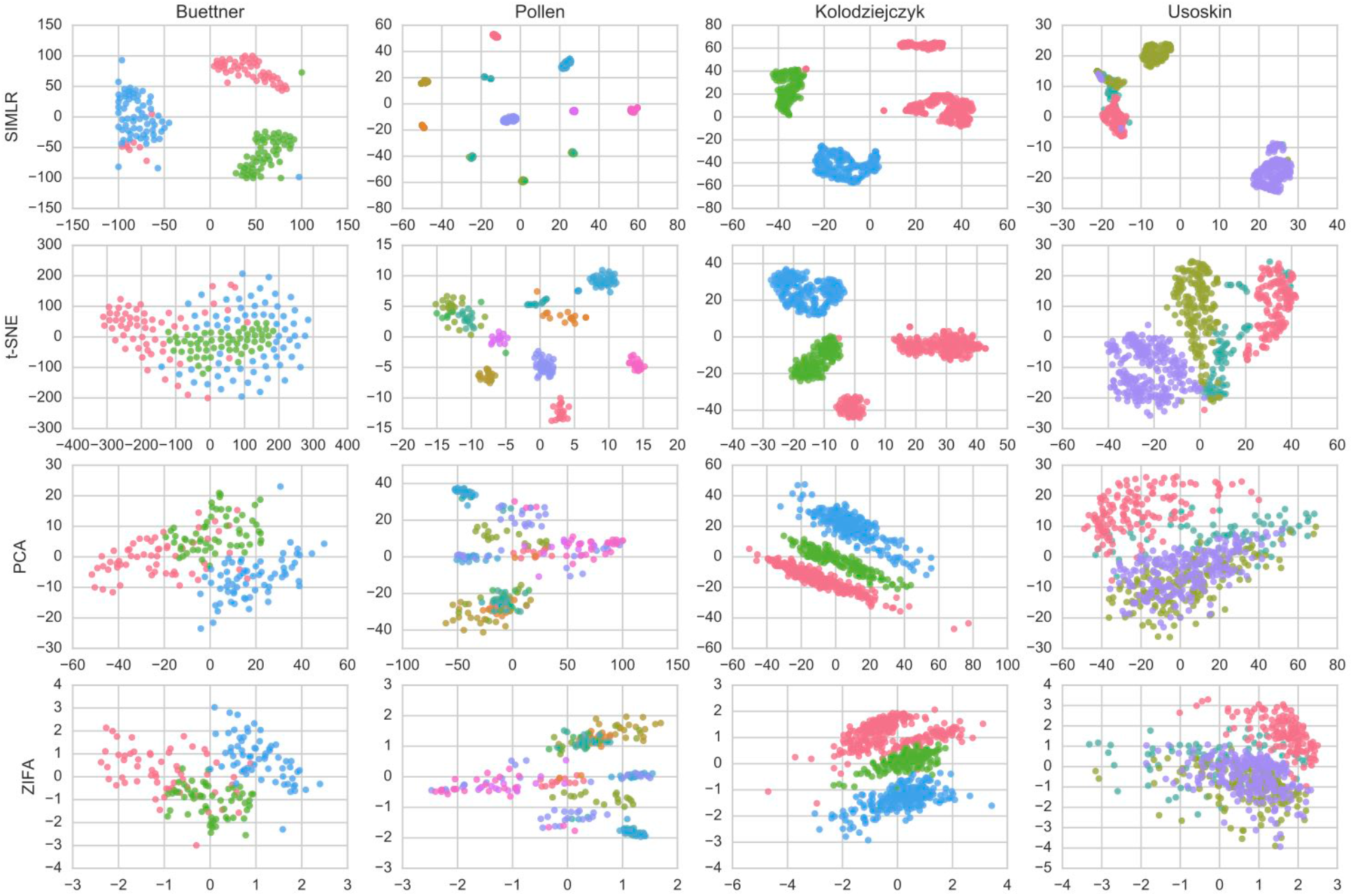
Comparison of different representative dimension reduction methods for visualizing four single-cell data sets. Each row corresponds to one of four different methods (SIMLR, t-SNE, PCA, and ZIFA) and each column corresponds to a different data set ([8]–[10], [17]). Each point is colored by its true label in the corresponding data set. There are 3 labels for the Buettner data set, 11 for the Pollen data set, 3 for the Kolodziejczyk data set, and 4 for the Usoskin data set.

SIMLR successfully separates the clusters for the Pollen (11 populations) and the Buettner (3 populations) data sets, whereas other methods contain clusters that are mixed to different extents. For the 3 populations in the Kolodziejczyk data set, SIMLR and t-SNE perform similarly and separate the clusters more clearly than PCA and ZIFA. For the 4 populations in Usoskin, none of the methods separated the clusters completely but SIMLR and t-SNE exhibit less overlap than PCA and ZIFA. We also performed additional comparisons between SIMLR, t-SNE, PCA and ZIFA on simulated data sets with ground-truth information to demonstrate SIMLR's superior performance under different modeling assumptions (**Supplementary Figures 7-13, 21-24 and Supplementary Note 3.4, 3.5, 3.6**).

These results indicate that SIMLR overall uncovers meaningful clusters that are more identifiable than those produced by existing methods. The visualizations of the four data sets using other dimension reduction methods are provided in **Supplementary Figure 1** for reference.

### Application to sparse PBMC data sets

Single-cell RNA-seq data produced from high-throughput microfluidics platforms such as DropSeq [13], InDrop [14] and GemCode single-cell technology [15] contains up to 95% zero expression counts. We tested the performance of SIMLR on sparse data sets by applying it to two PBMC scRNA-seq data sets from the GemCode [15]: PBMC3k and *in silico* PBMC (for the full details of these data sets, see **Materials and Methods**).

The *in silico* PBMC data set consists of 5 purified immune cell populations: naive *B* (CD19+ and IgD+), CD56+ natural killer (NK) cells, CD8+ cytotoxic T cells, CD4+ T cells and CD14+ monocytes. We generated *in silico mixtures* from these 5 data sets at specific proportions to generate ground-truth sets and SIMLR produced highly consistent unbiased classification of the true cell types (with NMI over 0.95 on average, **Supplementary Figures 7-8, Supplementary Note 3.2**).

In contrast to the *in silico* mixtures, the PBMC3k data set consists of the 5 major cell types and additional cell types in a healthy human. Because there is no ground-truth cell type information for the PBMC3k data set, we validated SIMLR's *de novo* cell type classification on the 2700 PBMCs using specific gene markers. We were able to identify the 5 major cell types as well as a rare megakaryocyte population of 12 cells with 0.44% abundance in the data set (**Supplementary Figure 16, Supplementary Table 4, and Supplementary Note 3.7.2**).

### Application to large scale datasets

To demonstrate the scalability of SIMLR, we applied it to three large scale single-cell data sets (for the full details of these data sets, see **Materials and Methods**):

1. Cells from the mouse cortex and hippocampus (Zeisel data set [39]).
2. Mouse retina cells with 39 subtypes (Macoskco data set [13]).
3. PBMCs from a healthy human (PBMC68k data sets [15]).

The analysis of these datasets consists of three components: (1) learning the cell-to-cell similarities with SIMLR, (2) grouping the cells by applying hierarchical clustering directly on the learned similarities, and (3) visualizing the cells in 2-D using the adaptive SNE framework with SIMLR.

The time and space complexity of these components (Table 5 and **Supplementary Table 3**) suggests SIMLR's feasibility on large-scale datasets. Because the cell types in each of the three data sets were originally assigned computationally, we use NMI as a measure of consistency between our classification labels and the original classification labels (Table 5). Moreover, we visualize the original classification labels from the studies using SIMLR's modified SNE framework to demonstrate SIMLR's effectiveness in revealing meaningful structures in its 2-D embedding (**Supplementary Figure 26**). Hierarchical application of SIMLR on Zeisel et al. indicates that SIMLR can dissect fine-grained heterogeneity in single-cell RNA-seq data (Table 5).

**Table 5.**
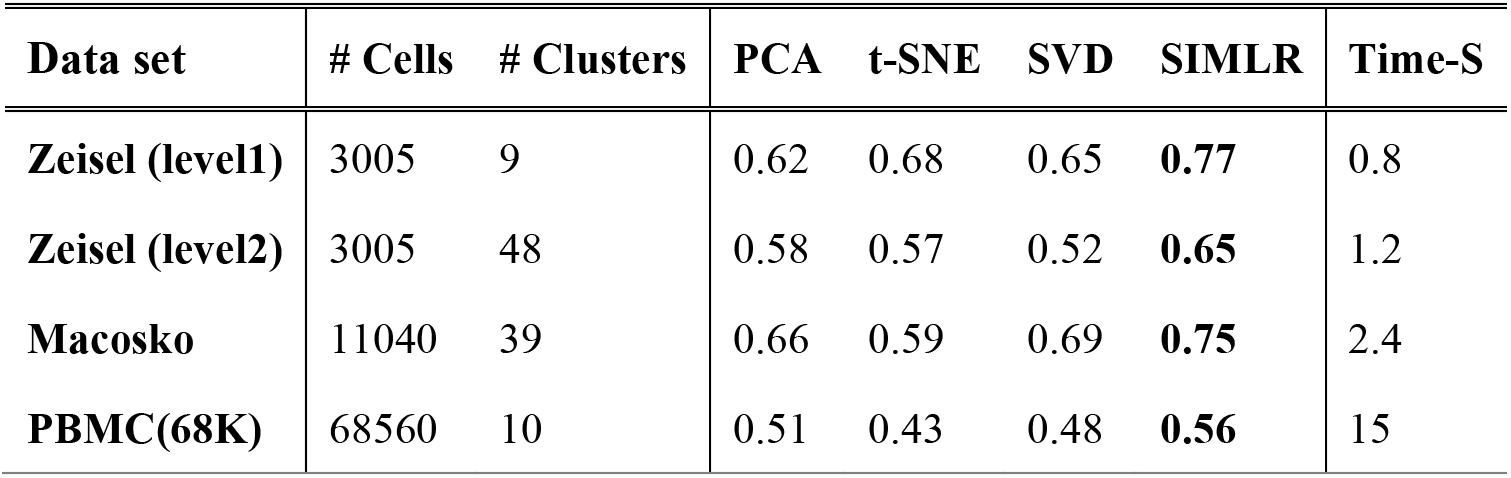
Summary and results on the three large-scale single-cell data. In Zeisel et al., we consider different levels of heterogeneity, which contain 9 and 40 sub-populations respectively. We report NMI values for four different methods. Last column shows the running time of SIMLR on these datasets. Time is reported in minutes.

## Discussion

High-throughput single-cell RNA sequencing technologies have enabled fine-grained analysis of cell-to-cell heterogeneity and molecular functions within tissues and cell populations that until recently could only be studied in bulk. As additional high-throughput approaches become available for single-cell RNA-seq, a wider range of studies pertaining to fundamental cellular functions will continue to emerge. Consequently, single-cell data sets may exhibit even higher levels of diversity (e.g., different tissues, comparisons of healthy versus disease-associated cell populations, different stages of the cell cycle or across development, different experimental technologies, and other sources of variation across data sets). Such diversity in experimental conditions and cell collections makes it difficult and undesirable to define cell-to-cell similarity measures based on strict statistical assumptions. As novel methodologies and algorithms are urgently needed for this new type of data, SIMLR aims to adapt to the heterogeneity across different single-cell data sets by learning an appropriate similarity measure for each data set.

In this paper we extensively evaluated SIMLR using recently published distinct single-cell data sets and without any prior knowledge of groups of significant genes. We demonstrated that SIMLR is able to learn appropriate cell-to-cell distances that uncover similarity structures that would otherwise be concealed by noise or outlier effects. Furthermore, SIMLR successfully clusters cell populations and projects the high-dimensional data in a visually intuitive fashion. We show that SIMLR separates clusters more cleanly than 8 other popular dimension reduction methods, including linear methods (such as PCA), nonlinear methods (such as Sammon), and a recently published approach (ZIFA) specifically designed for single-cell data sets [12].

Because each dimension reduction algorithm makes its own assumptions, it is unlikely that one is optimal for all data sets. As we have shown, SIMLR performs well on single-cell data sets that contain several clusters - a frequent use case where heterogeneity is defined by distinct cell lineages. But because the current implementation of SIMLR assumes the data has cluster structures, it may not be best suited for data that does not contain clear clusters, such as cell populations that contain cells spanning a continuum (such as progression through pseudotime [40], [41]). Methods that construct this continuum also rely on cell-to-cell similarities which contain noise and high levels of sparsity, and a multiple-kernel learning framework may still be beneficial. Therefore, it will be interesting to develop approaches like SIMLR customized for such data sets which do not contain clear clusters.

## Materials and Methods

### Four Published Data Sets

We used the following four data sets in our analysis:

1. Eleven cell populations including neural cells and blood cells (Pollen data set [9]). This data set was designed to test the utility of low-coverage single-cell RNA-seq in identifying distinct cell populations, and thus contained a mixture of diverse cell types: skin cells, pluripotent stem cells, blood cells, and neural cells. This data set includes samples sequenced at both high and low depth; we analysed the high-depth samples, which were sequenced to an average of 8.9 million reads per cell.
2. Neuronal cells with sensory subtypes (Usoskin data set [8]). This data set contains 622 cells from the mouse dorsal root ganglion, with an average of 1.14 million reads per cell. The authors divided the cells into four neuronal types: peptidergic nociceptors, non-peptidergic nociceptors, neurofilament containing, and tyrosine hydroxylase containing.
3. Embryonic stem cells under different cell cycle stages (Buettner data set [17]). This data set was obtained from a controlled study that quantified the effect of the cell cycle on gene expression level in individual mouse embryonic stem cells (mESCs). An average of 0.5 million reads were obtained for each of the 182 cells and at least 20% of the reads were mapped to known exons on the mm9 mouse genome. The cells were sorted for three stages of the cell cycle using fluorescence-activated cell sorting, and were validated using gold-standard Hoechst staining.
4. Pluripotent cells under different environment conditions (Kolodziejczyk data set [10]). This data set was obtained from a stem cell study on how different culture conditions influence pluripotent states of mESCs. This study quantified the expression levels of about 10 thousand genes across 704 mESCs from 9 different experiments involving three different culture conditions. An average of 9 million reads were obtained for each cell and over 60% of the reads mapped to exons on the *Mus musculus* genome.

For all the data sets above, we applied a logarithmic transformation *f*(*X*) = log_10_(*X* + 1) to the single-cell raw expression data.

### Three Large-scale Data Sets

1. Cells from the mouse cortex and hippocampus (Zeisel data set [39]), collected using unique molecule identifier (UMI) assays and 3'-end counting. 3,005 cells from the mouse brain were collected, and 47 sub-types were identified by hierarchical bi-clustering and validated by gene markers.
2. Mouse retina cells with 39 subtypes (Macoskco data set [13]). Obtained by Drop-seq, a droplet-based high-throughput technique[13], this data set includes UMI (3'-end) counts for 44,808 cells (identified by their customized computational pipeline). The cell types were classified via PCA and density-based clustering, and validated by differential gene expression. Similar to the processing procedure used in [13], we filtered out cells with less than 900 genes (resulting in 11040 cells) for unsupervised analysis.
3. PBMCs from a healthy human (PBMC3k and PBMC68k data set [15]). scRNA-seq libraries were generated by 10x Genomics GemCode platform, a droplet-based high-throughput technique, [15] and 2700 cells with UMI (3'-end) counts were identified by their customized computational pipeline. This cell population includes major immune cell types in a healthy human.

### Multiple Kernel Learning

Instead of using a predefined distance metric, we incorporate multiple kernel learning in SIMLR to compute the distances between pairs of cells. The general form of the distance between cell *i* and cell *j* is defined as

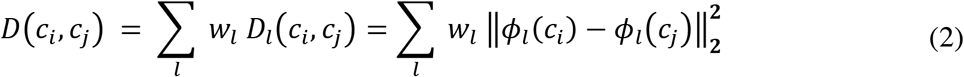

where *ϕ_l_*(*c_i_*) is the *l*th kernel-induced *implicit* mapping of the *i*th cell. This mapping is implicit because we are only concerned about the inner products of the *ϕ_l_*(*c_i_*) and *ϕ_l_*(*c_j_*) for pairs (*i,j*) which is defined as follows:

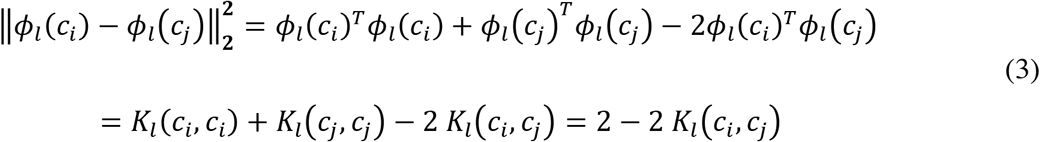

where we only need to compute the kernel functions *K_l_*(*c_i_*, *c_j_*). (The kernel of two identical inputs is set to 1 by convention). So we can refine the optimization problem to include the distance in the objective and the corresponding weights as variables as follows:

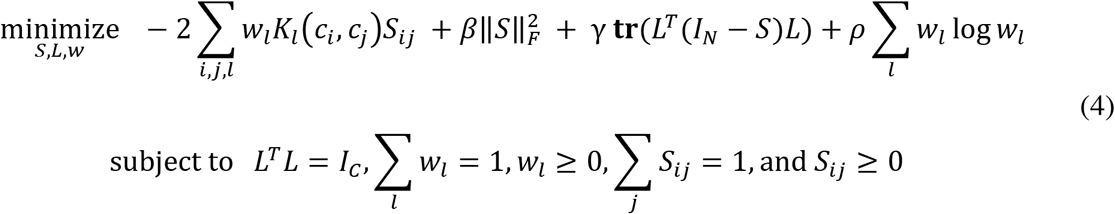

We describe the optimization of *L, w*, and *S* below; we describe selection of the parameters *β*, *γ* and *ρ* in the **Supplementary Note 3.8** and **3.11**.

### Kernel Construction

In the default implementation of SIMLR, we use Gaussian kernels with various hyper-parameters. Gaussian kernels, which generate superior empirical performances than other types of kernels, take the form

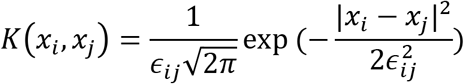

where *x_i_* and *x_j_* denote the i-th and j-th cell respectively. The variance *ϵ_ij_* can be calculated with different scales:

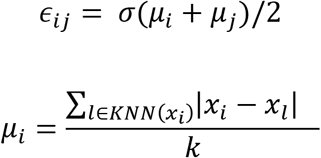

where *KNN*(*x_i_*) represents cells that are top k neighbors of the i-th cell. Hence, each kernel is decided by a pair of parameters (*σ*,*k*). We set *k* = 10,12,14,…, 30 and *σ* = 1.0,1.25,1.5,1.75,2, resulting in 55 different kernels in total. However, our method is relatively insensitive to the number of kernels and choices of parameters (**Supplementary Figure 28**).

### Solving the Optimization Problem

Initialization of *S, w*, and *L*: The weight of multiple kernels, *w*, is initialized as an uniform distribution vector, i.e., 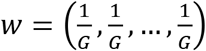, where *G* is the number of kernels. The similarity matrix *S* is initialized as *S_ij_* = ∑_l_ *w_l_ K_l_*, (*c_i_, c_i_*). *L* is initialized as the top *C* eigenvectors of *I_N_* − *S*.

The optimization problem formulated above is non-convex with respect to all of the variables *S, L, w*, but the problem of each variable conditional on other variables being fixed is convex. So we can apply an alternating convex optimization method to solve this tri-convex problem efficiently. The following four steps listed in Algorithm (1) are implemented iteratively.

#### Step 1: Fixing L and w to update S

When we minimize the objective function with respect to (w.r.t.) the similarity matrix *S*, we can rewrite the optimization problem as follows.

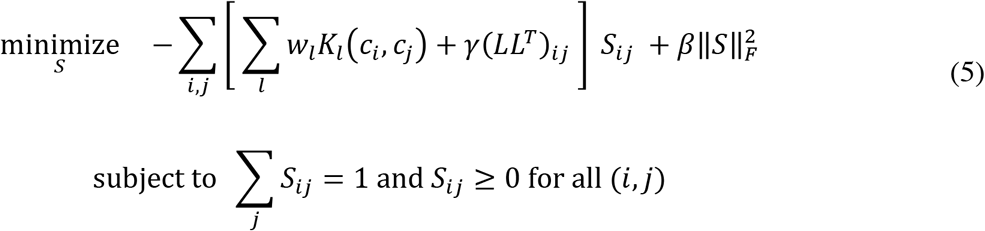

The first summation term in the objective as well as constraints are all linear, and the second summation in the objective is a simple quadratic form that can be solved in polynomial time [42]. We provide details on how SIMLR is implemented to solve this problem efficiently in **Supplementary Note 3.8.1.**

#### Step 2: Fixing S and w to update L

When we minimize the objective function w.r.t. the latent matrix *L*, we can rewrite the optimization problem as follows.

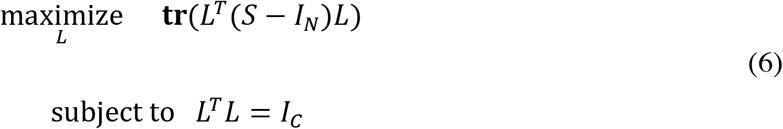

The trace of *L^T^* (*S* − *I_N_*)*L* is maximized when *L* is an orthogonal basis of the eigenspace associated with the *C* largest eigenvalues of (*S* – *I_N_*) [43]. Thus, *L* can be computed efficiently using any matrix numeric toolbox (with implementation details provided in **Supplementary Note 3.8.2**).

#### Step 3: Fixing S and L to update w

When we minimize the objective function w.r.t. the kernel weights *w*, we can re-write the optimization problem as follows.

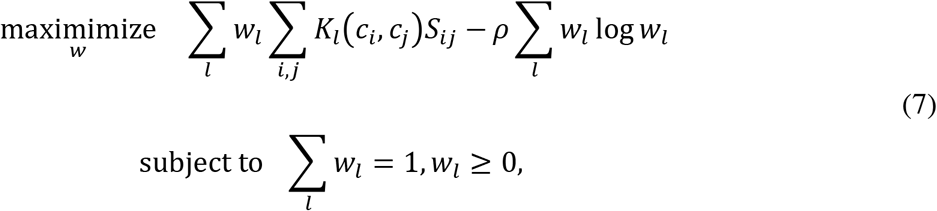

The problem with a convex objective and linear constraints can be solved by any standard convex optimization method [42]. Details of the step of updating wand choices of the specific multiple kernels we choose are stated in **Supplementary Note 3.8.3**.

#### Step 4: Similarity enhancement

We apply a diffusion-based step to enhance the similarity matrix *S* and reduce the effects of noise and especially dropouts in single-cell data. Given *S*, we construct a transition matrix *P* such that

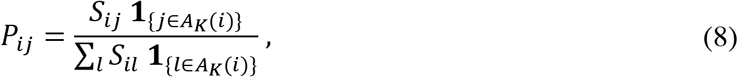

where *A_K_* (*i*) represents the set of indices of cells that are the *K* top neighbors of cell *i* under the learned distance metric. Under this construction, the transition matrix is sparse so we preserve most of the similarity structure. The diffusion-based method that we apply to enhance the similarity *S* has the following update scheme:

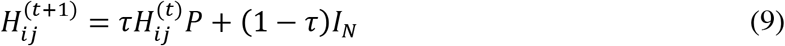

where we have 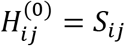 as an input and the final iteration of *H_ij_* is used as the new similarity measure *S_ij_*-. We show that the diffusion process converges in **Supplementary Note 3.8.6**.

SIMLR iterates the four steps above until convergence. We use the eigengap, defined as the difference between the *C* + 1th and *C*th eigenvalues [44], (**Supplementary Note 3.8.5**) as the convergence criterion. When the method has converged, the similarity *S* should be stable and so its eigengap should be stable too. Further, a good low rank similarity *S* should have a small eigengap. We show the dynamics of the eigengap during iterations in SIMLR on the four real data sets (**Supplementary Figure 18**). SIMLR converges within around 10 iterations.

### Large-Scale Extension of SIMLR

We extend SIMLR to deal with large scale single-cell RNA-seq datasets of tens of thousands of cells. The key idea is to use K-nearest-neighbor (KNN) similarity to approximate the full pair-wise similarity. The first step is to use a state-of-art nearest neighbor search technique kGraph [45]. kGraph optimizes indexing of KNN graph by relying on the heuristic that a neighbor's neighbor is probably a neighbor too. After the construction of KNN graph, we only update similarities in these pre-selected top K neighbors for each cell. Since the similarity is sparse, we use a representative eigen-decomposition library named Spectra [46] to solve for *L* in Eqn.(6). Detailed optimization procedures in solving variables *S* and *L* are provided in **Supplementary Note 3.12**. As to similarity enhancement, instead of calculating the closed-form solution which involves a large matrix inversion, we only apply limited number of iterations to get an approximation of the final solution.

After we obtain the cell-to-cell similarity, we can perform both cell subpopulation identification and cell visualization. When clustering the cells in large scale, it is computationally expensive to obtain embedding from tSNE-type framework. Instead, we modified a hierarchical clustering algorithm for similarity which hierarchically merges small clusters into big cluster based on similarity density until a desired number of clusters is achieved. This simple algorithm is very effective in sparse similarity and scales up to tens of thousands of cells without explicitly calculating the embedding.

For visualization, since we are only mapping cell-to-cell similarity into 2D or 3D space, it is still computationally tractable to apply a tSNE-type embedding. However, we modified the famous Barnes-Hut algorithm in tSNE [47] by utilizing the sparse properties of our similarity (detailed in **Supplementary Note 3.12.3**).

## Software Availability

SIMLR is freely available as both a MATLAB program and an R package (https://github.com/BatzoglouLabSU/SIMLR). For K-means clustering, we used the kmeans implementation for MATLAB [48] and the built-in function in R. For stochastic neighbor embedding, we modified the source code of the t-SNE implementation in MATLAB [19] and that in R [49] respectively.

## Additional Files

The following additional file is available with this paper. Additional file 1: **Supplementary Information** includes Supplementary Tables, Figures and Notes.

## Abbreviations

SIMLR: Single-cell Interpretation via multi-kernel enhanced similarity learning
mESCs: mouse embryonic stem cells
NMI: Normalized mutual information
NNE: Nearest neighbor error
MDS: Multidimensional scaling
FA: Factor analysis
PCA: Principal component analysis
PPCA: Probabilistic principal components analysis
ZIFA: Zero-inflated factor analysis
SNE: Stochastic neighbor embedding
t-SNE: t-distributed stochastic neighbor embedding

## Authors' contributions

BW, JZ and SB conceived the study and planned experiments. BW designed the algorithm and implemented the software in MATLAB. DR developed the software package in R. BW, JZ and EP performed data analysis implemented the simulation study. JZ and EP drafted the manuscript. BW and SB contributed to the manuscript. All authors read and approved the final manuscript.

## Acknowledgements

The authors would like to thank Grace X Zheng, Jessica Terry, Tarjei Mikkelsen from 10x Genomics for providing access to the PBMC data as well as suggestions for the manuscript and the *in silico* experiments. EP acknowledges support from an NDSEG Fellowship and a Hertz Fellowship. JZ acknowledges support from Stanford Graduate Fellowship.

## Algorithms

**Algorithm 1.**
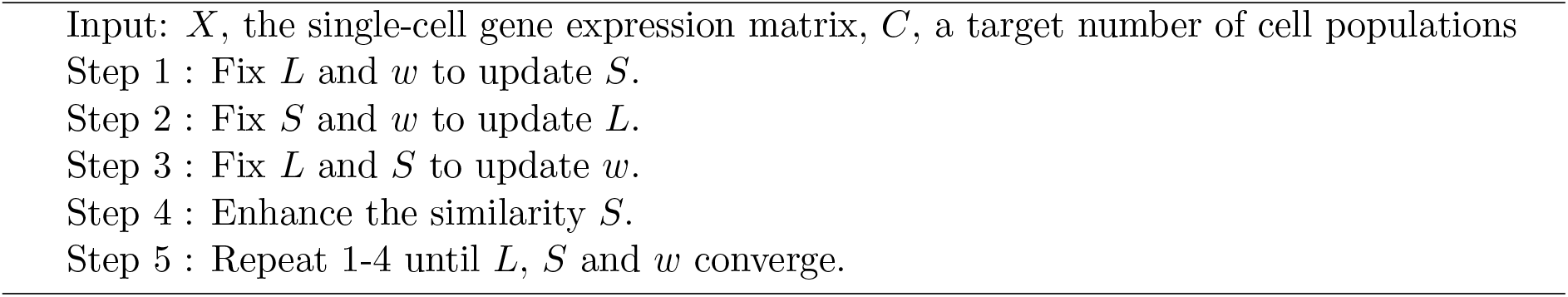
Similarity Learning via SIMLR.

